# Myc-dependent cell competition and proliferative response requires induction of the ribosome biogenesis regulator Peter Pan

**DOI:** 10.1101/2020.05.06.080283

**Authors:** Norman Zielke, Anna Vähärautio, Jianping Liu, Jussi Taipale

**Affiliations:** Applied Tumor Genomics Research Program, University of Helsinki, Finland; Research Program in Systems Oncology, University of Helsinki, Finland; Department of Medical Biochemistry and Biophysics, Karolinska Institutet, Stockholm, Sweden; Department of Biochemistry, University of Cambridge, UK

## Abstract

The transcription factor Myc is activated in most major forms of human cancer. Myc regulates a large set of target genes, and drives cell growth across animal phyla. However, it has not been clear which target genes are required for Myc-induced growth, and whether the targets are individually necessary or act in an additive fashion. Here, we have used comparative functional genomics to identify a core set of Myc target genes whose regulation is conserved between humans and *Drosophila melanogaster*. Most of these targets are essential genes involved in ribosome biogenesis and ribonucleotide metabolism. To identify *Drosophila* genes whose upregulation is necessary for Myc induced growth, we deleted the Myc binding sites (E-boxes) in the promoter regions of four genes using CRISPR/Cas9. All mutant flies were homozygous viable, indicating that E-box sequences are not required for basal expression of the Myc target genes. E-Box deletions in RpS20, RpS24 and Nop56 did not cause strong growth phenotypes. However, deletion of the E-box in the rRNA processing factor *Peter Pan* (*ppan*) made the flies resistant to Myc-induced cell growth, without affecting Myc-induced apoptosis. Despite their failure to respond to Myc, the *ppan^Ebox−/−^* flies are healthy and display only a minor developmental delay, suggesting that it may be possible to treat or prevent tumorigenesis by targeting individual downstream targets of Myc.

## Main Text

The transcription factor Myc is commonly activated by upstream oncogenic pathways and amplification of the Myc locus is one of the most common genetic alterations in cancer genomes (*1*, *2*). Myc functions mainly as a transcriptional activator that binds to DNA in conjunction with its obligate heterodimeric partner Max. Both Max:Max homodimer and Myc:Max heterodimer can bind with high affinity to sites containing the sequence CACGTG (E-box) or its close variants (*3*–*7*). Transient expression of a genetically encoded Myc inhibitor (Omomyc) impairs malignant growth (*8*, *9*), qualifying Myc as a prime target for the development of anti-cancer therapeutics. Furthermore, loss of Myc regulatory regions makes mice resistant to cancer (*10*, *11*), suggesting that inhibition of Myc could also be used in cancer chemoprevention. However, the Myc:Max heterodimer lacks binding pockets that would facilitate binding of small molecule drugs (*12*), making it difficult to inhibit the activity of the Myc protein pharmacologically. Identifying crucial downstream targets of Myc is therefore a promising alternative route, but this approach is complicated by the large number of Myc target genes, many of which are regulated in a tissue- or cell type specific manner (*7*, *13*–*17*). Furthermore, Myc has been proposed to act as a universal transcriptional amplifier that boosts the expression of all genes (*7*, *13*), making it unclear whether its activity could be blocked by interfering with the induction of a single target gene.

We reasoned that because different cell types respond to Myc similarly by inducing growth, the target genes that drive growth should be activated by Myc in multiple different cell types. Furthermore, as the phenotypic effects of Myc are largely conserved between animal species, the central downstream mechanisms driving growth are also likely to be conserved. Therefore, to address the role of Myc target genes in cell growth regulation, we set out to identify conserved target genes that are regulated by Myc irrespective of species or cell type. As a model system, we chose *Drosophila*, as the fly Myc gene (dMyc) is functionally equivalent to mammalian Myc (*18*, *19*), and can transform rodent fibroblasts together with H-Ras (*20*). Furthermore, *Drosophila* provides an exceptionally powerful *in vivo* genetic toolset to study Myc activity, and is sufficiently far in evolutionary distance from humans to limit the number of functionally conserved targets.

To determine targets of Myc that are functionally conserved between humans and *Drosophila*, we used RNAi followed by expression profiling (RNA-seq) and chromatin immunoprecipitation followed by sequencing (ChIP-seq) in two human colorectal cancer cell lines (GP5d, LoVo) and in *Drosophila* hemocyte-derived S2 cells. Targets were identified using relatively loose criteria to avoid false negatives (see **Materials and Methods**), resulting in a list of 124 *Drosophila* genes, that correspond to 126 orthologous genes in humans (**Figure 1A**). The resulting list of functionally conserved targets **(Table S1)**is partially overlapping with prior studies, which have identified conserved target sets for Myc based on experimental data and/or conservation of E-box motifs in promoters (*5*, *6*, *16*, *21*). Consistent with co-evolution of Myc with a ribosome biogenesis regulon (*22*, *23*), the functionally conserved targets were highly enriched in ribosomal components, and in regulators of ribosome biogenesis and ribonucleotide metabolism (**Figure 1A**). Interestingly, almost all conserved target genes in *Drosophila* (114 of 124 genes with positive fold change in wing disc after dMyc overexpression) we identified were upregulated by Myc (**Figure 1B**), suggesting that the conserved function of Myc is to act as a transcriptional activator.

**Figure 1.**
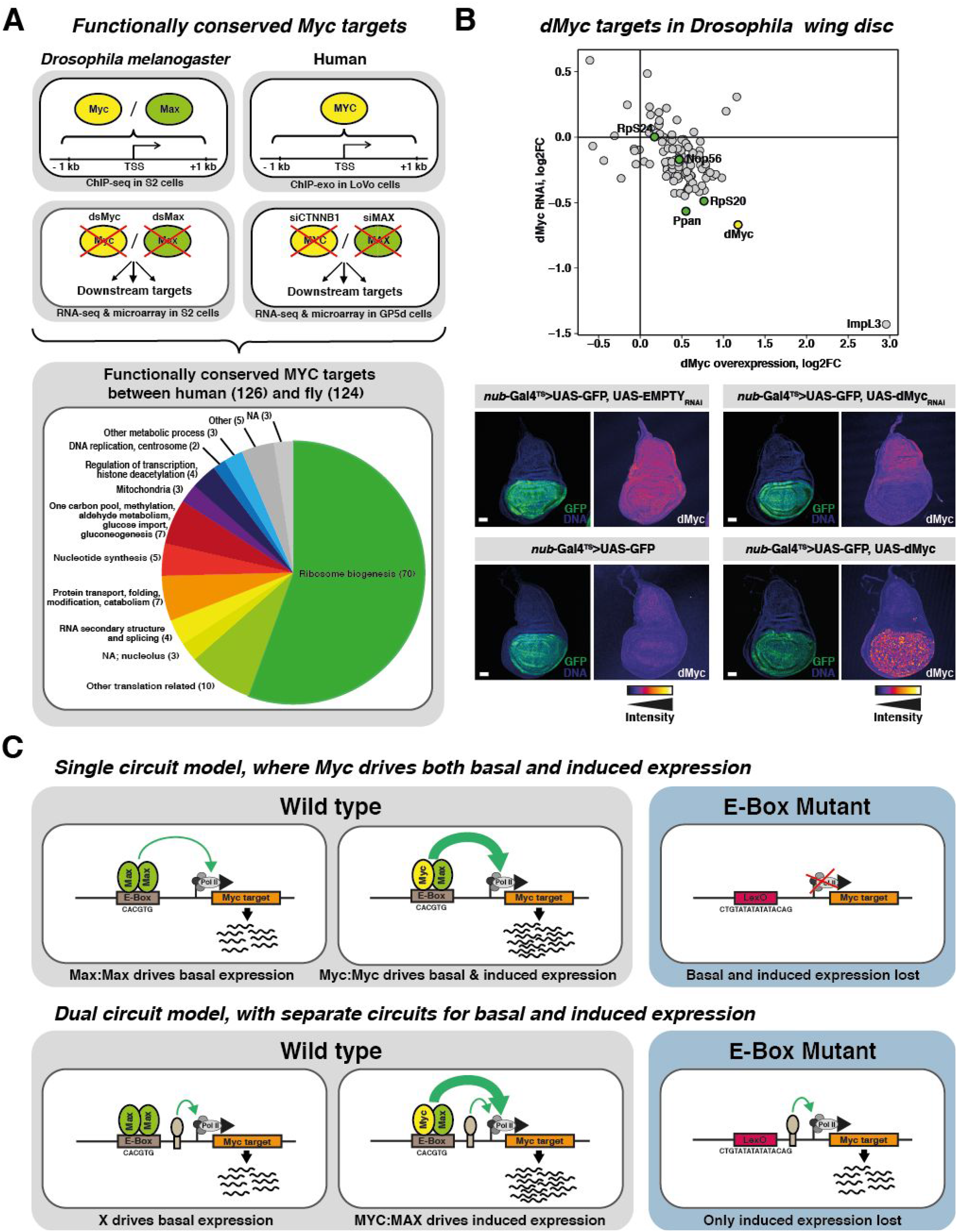
Identification of a functionally conserved set of MYC/Max target genes. **(A)** Overview of the strategy for the identification of functionally conserved Myc target genes between humans and flies. Analysis was based on the presence of ChIP-seq peaks (top panels) and change in expression after downregulation of Myc/Max activity (middle panels). Pie chart (bottom panel) shows classification of target genes according to human Biological Process Gene Ontology (GO) terms. **(B)** Effect of dMyc overexpression (x-axis) or RNAi repression (y-axis) on conserved Myc targets in the *Drosophila* wing disc (upper panel). Lower panel shows confocal micrographs of wing imaginal discs in which either UAS-Myc or UAS-MycRNAi as well as the respective controls were conditionally overexpressed in the wing pouch using *nub*-Gal4; tub-Gal80^TS^. DNA is stained with Hoechst 33342 and shown in blue, GFP is shown in green. The color scale of dMyc protein expression is shown below the pictures. Scale bars: 50 *μ*m. **(C)** Schematic representation of the regulation of the Myc/Max target genes. Please note that in the absence of Myc activity, the basal level of expression of the targets could be driven by Max or other E-box binding TFs (top right), or alternatively be regulated independently of the E-box by other constitutively active TFs (top left). Therefore, deletion of the E-boxes could either lead to a complete loss of expression (null phenotype, bottom left), or retention of basal expression and specific loss of regulation of the targets by Myc (bottom right).

As most of the functionally conserved Myc target genes were known essential genes, and had a basal expression level even after RNAi targeting of Myc, a straightforward genetic dissection of the regulatory network activated by Myc was not feasible. This is because rescue-experiments are extremely challenging, as a large number of target genes would have to be overexpressed to rescue a Myc loss-of-function phenotype (*24*, *25*). In addition, due to the basal expression, loss-of-function mutations of downstream components will have a phenotype that is distinct from mutations that specifically prevent their regulation by Myc (see for example Ref. (*26*)). Since mutation of E-boxes has previously been shown to abolish the regulation by Myc in reporter constructs (*27*),(*27*)we decided to directly replace the E-boxes in a set of conserved Myc target genes with LexO sites in using CRISPR/Cas9 mediated gene editing **(Figure S1)**. One potential outcome of this experiment would be that genomic deletion of the E-boxes uncouples basal and Myc-induced expression of the target genes. Another possibility is that E-boxes are required for basal expression, as the Max:Max homodimer, and many other bHLH proteins can also bind to E-boxes (*28*–*30*)e latter possibility would result in reduced viability when E-boxes of essential target genes are mutated (**Figure 1C**).

For our analysis, we selected the ribosomal proteins RpS20 and RpS24 (*31*), and two ribosome biogenesis regulators, the U3 snoRNP component Nop56 (*32*) and *Peter Pan* (*ppan*), the bona fide ortholog of yeast SSF1 and 2, which are involved in the processing of the 27SA2 pre-rRNA (*33*, *34*). These genes were selected because 1) they are all essential genes with expected growth phenotypes, 2) they have both dMyc and dMax ChIP-seq peaks at their promoters that overlap consensus E-boxes, and 3) the E-boxes have nearby GG protospacer adjacent motif (PAM) sites that facilitate CRISPR/Cas9-mediated gene editing **(Figure S1)**.

The mutant lines were generated using CRISPR/Cas9, and subsequently bred to homozygosity (see **Materials and Methods**). All of them (*RpS20^Ebox−/−^*, *RpS24^Ebox−/−^*, *Nop56^Ebox−/−^* and *ppan^Ebox−/−^*) were viable, indicating that the deleted E-boxes were not critically important for the basal expression of these essential genes. We next determined whether the homozygous E-box null flies displayed a growth phenotype. Many ribosomal proteins and ribosome biogenesis factors belong to the class of *Minute* genes, haploinsufficient loci that delay cellular growth and larval development, but give rise to almost normal-sized flies with a subtle, characteristic thin bristle phenotype (*31*, *35*) **(Figure 2A)**. We used scanning electron microscopy followed by quantitative imaging to measure the scutellar bristle lengths of the homozygous lines (**Figure 2B**; n > 10). This analysis revealed that there was no measurable difference between *RpS24^Ebox−/−^* and w^1118^ control flies (p = 0.169; Welch’s T-test) whereas *RpS20^Ebox−/−^* (p < 0.0001) displayed a small decrease in bristle length compared to control flies, which was less prominent than the phenotype observed in heterozygotes of the known *Minute* mutation *RpS24*^SH2053^ (p < 0.0001). *Nop56^Ebox−/−^* showed an intermediate phenotype, with moderately decreased bristle length compared to control flies (p < 0.0001) that was significantly different from the prominent short bristle phenotype observed in heterozygotes of *RpS24*^SH2053^ (p = 0.0174). In contrast, homozygous loss of the E-box in *ppan* resulted in a strong bristle phenotype (p < 0.0001) that was indistinguishable from the *RpS24*^SH2053^ heterozygotes (p = 0.0695). Consistently, analysis of developmental timing revealed that the larval development of *ppan^Ebox−/−^* flies was delayed; the onset of pupariation was similar to that seen in the *Minute* flies (RpS24^SH2053^; **Figure 2C**). The effect on development to adulthood was even more severely delayed (**Figure 2D**). Interestingly, the effect was more prominent in males than in females, potentially because of sex differences in the expression of the Myc gene (*36*), which is located in the X chromosome in flies. Importantly, most known *Minute* mutations, except Myc, are haploinsufficient and homozygous lethal (*31*). Similar to *Myc^P0 (37)^*, whose phenotype is only detected in the hemizygous state, the *Minute-like* phenotype of *ppan^Ebox−/−^* is only observable in a homozygous state, providing strong genetic evidence for *ppan* being a critical downstream mediator of Myc.

**Figure 2.**
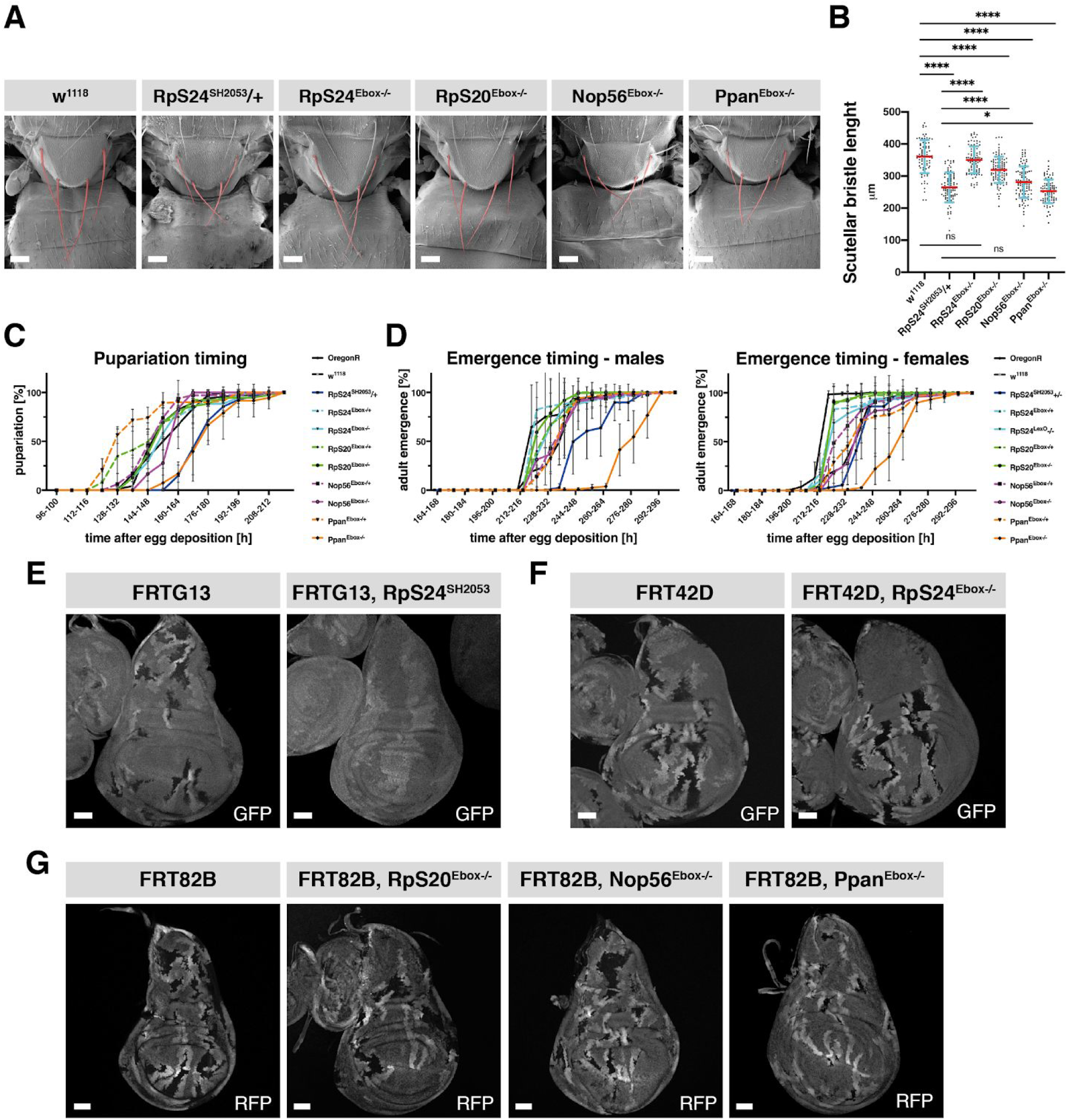
E-box of Ppan is required for Myc dependent growth. **(A)** Scanning electron micrographs of *Drosophila* scutella showing the characteristic bristle phenotype. Heterozygotes of the known *Minute* mutation *RpS24*^SH2053^ were included as controls. Scutellar bristles are indicated in red. Scale bars: 100 *μ*m. **(B)** Quantification of scutellar bristle lengths from (A). Mean values are represented by a red bar. Error bars indicate standard deviation (SD). **(C)** Pupariation timing of E-box null larvae (orange) compared to wild-type (OregonR, black) and *Minute* files (blue). Fraction of emerged pupae (y-axis) as a function of time after egg deposition (x-axis) is shown. Genotypes are indicated in the legend. Error bars indicate SD. **(D)** Comparison of emergence time of adults. Fractions of the emerged male (left panel) and female (right panel) adult flies (y-axis) as a function of time after egg deposition (x-axis) is shown. . Genotypes are as in (C). Error bars indicate SD. **E-G)** Twin-spot analysis in *Drosophila* wing discs. Twin clones were generated using the heat-shock controlled Flp/FRT systems. Genotype and the used fluorescence reporter are indicated above and on bottom right of each panel, respectively. Homozygous clones are indicated by the absence of GFP or RFP. Scale bars: 50 *μ*m. E) Comparison of growth rates between FRTG13 and the known *Minute* mutation RpS24^SH2053^. F) Comparison of growth rates between FRT42D and RpS24^Ebox−/−^. G) Comparison of growth rates between FRT82B, RpS20^Ebox−/−^, Nop56^Ebox−/−^ and Ppan^Ebox−/−^ clones.

To determine whether regulation of the target genes is required for Myc-driven cell growth, we compared the growth-rates of the E-box^+/+^ and E-box^−/−^ clones in *Drosophila* wing imaginal discs. It has been previously shown that hypomorphic dMyc (*Myc^P0^*) clones as well as clones carrying *Minute* mutations or the *ppan* null allele (*ppan^j6B6^*) are eliminated by cell competition when surrounded by wild-type cells (*33*, *37*, *38*). As expected, we found that homozygous clones for a *Minute* allele (*RpS24^SH2053^*) became eliminated from wing discs, while the corresponding wild-type twin-clones increased in size compared to wild-type clones in control experiments **(Figure 2E)**. By contrast, homozygous clones for *RpS24^Ebox−^* and wild-type clones grew to sizes comparable to each other, indicating that *RpS24^Ebox−^* does not have measurable effect on cell competition or clone growth **(Figure 2F)**. Similarly, we observed no cell competition phenotypes in *RpS20^Ebox−^* or *Nop56^Ebox−^* flies **(Figure 2G)**, consistent with a model where the basal expression of these genes is sufficient for wing disc growth. By contrast, clones homozygous for the *ppan^Ebox−^* allele were at a clear growth disadvantage compared to wild-type clones, and were outcompeted in the wing disc **(Figure 2G)**, suggesting that wing disc cells are sensitive to small changes in *ppan* activity. These results show that regulation of *ppan* by endogenous levels of Myc is important for the growth of cells in the *Drosophila* wing imaginal discs. Taken together, these results indicate that regulation of only a subset of Myc target genes is rate limiting for growth, and suggest that regulation of rRNA precursor processing is of greater importance for Myc-induced growth than the transcriptional upregulation of individual ribosomal proteins.

Physiological levels of Myc are important for growth during development (*19*), whereas in tumors, Myc is often overexpressed (*1*). In *Drosophila* and in mice, overexpression of Myc converts cells into super-competitors that can outcompete adjacent wild-type cells (*39*–*42*). It has been proposed that super-competition could be involved in the premalignant stages of cancer, allowing mutant but phenotypically almost normal cells to outcompete wild-type cells, generating a field of pre-malignant cells (*43*, *44*). Moreover, apoptotic patterns characteristic for super competition were recently detected in the tumour-stroma interface and within the tumour parenchyma of human cancers, suggesting that super-competition could also be implicated in tumor invasiveness and metastasis (*45*). To address the role of the conserved Myc target genes in responding to supraphysiological levels of Myc, we used mosaic analysis with a repressible cell marker (MARCM) (*46*) to generate *ppan^Ebox−/−^* clones that overexpress Myc either alone, or together with the anti-apoptotic protein p35 in *Drosophila* wing discs. Staining of the wing disc using antibodies to a cleaved form of *Drosophila Death caspase 1* (Dcp1) revealed that Myc overexpression increased apoptosis within the clones and in cells directly adjacent to them (**Figure 3**). However, quantification of the clone sizes relative to the total area of the wing discs indicated that Myc overexpression did not significantly alter the size of the clones compared to wild-type (p = 0.31; one-sided Mann-Whitney U test). These results could be explained by induction of both cell proliferation and apoptosis by Myc (*47*). Consistently with this interpretation, co-expression of the p35 anti-apoptotic protein with Myc increased clone size significantly (p < 1.21 × 10^−11^) over clones expressing p35 alone (**Figure 3**). Deletion of the E-box of *ppan* prevented the increase in clone size induced by Myc and p35 co-expression, but did not affect Myc-induced apoptosis **(Figure 3)**. In summary, both the sister-clone (**Figure 2**) and MARCM (**Figure 3**) analyses indicate that Myc-driven proliferation requires the E-box of *ppan*, suggesting that limiting the activity of *ppan* could be a potent strategy for the treatment of Myc-dependent cancers.

**Figure 3.**
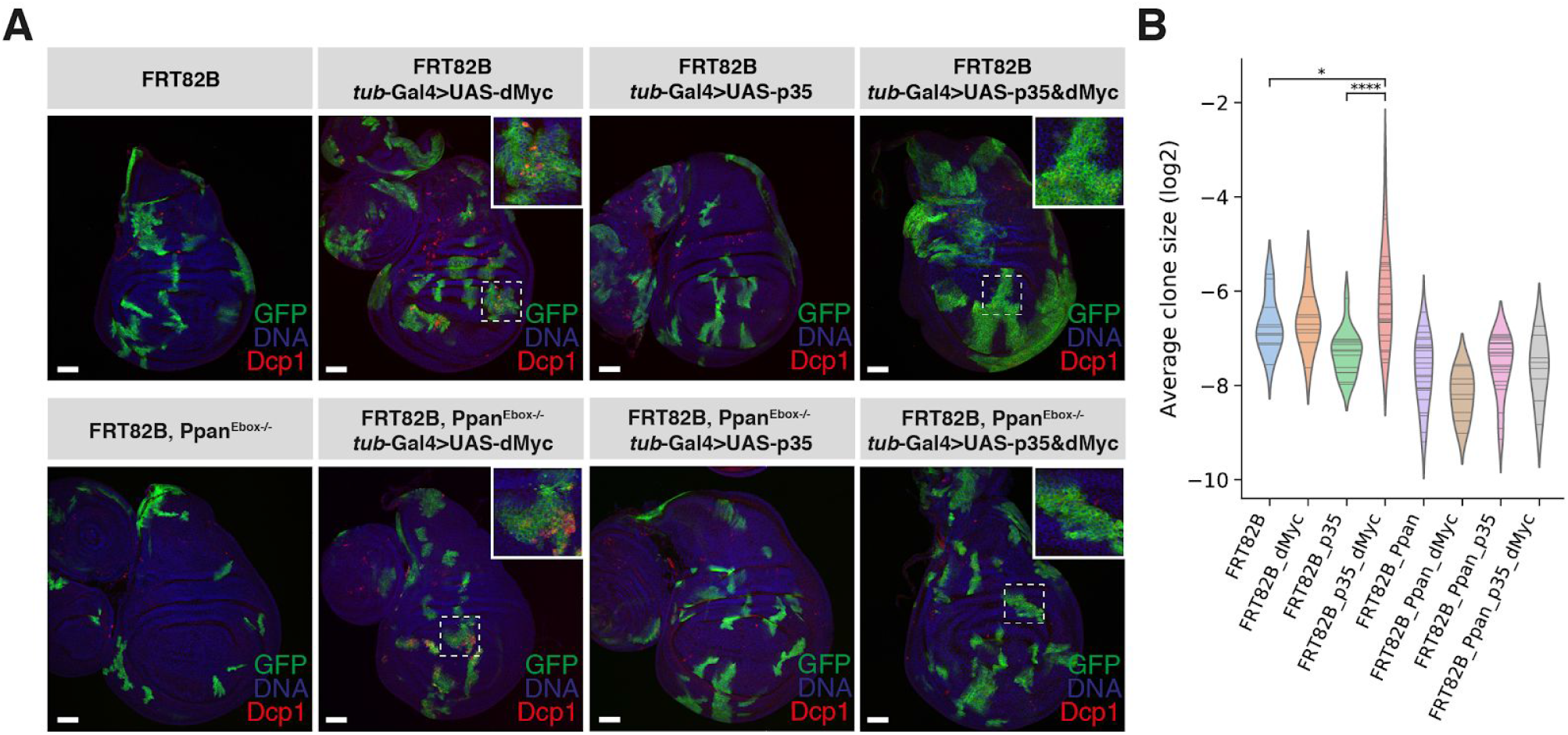
Deletion of an E-box in the Ppan promoter blocks responses to Myc. **A)**MARCM analysis of apoptosis and clone growth. Induced clones are marked with GFP. Note that dMyc overexpression increases apoptosis (Dcp1 staining) that is prevented by p35 but not loss of the Ppan E-box, and that loss of the Ppan E-box prevents clone growth induced by overexpression of dMyc together with p35. DNA is stained with Hoechst 33342 and shown in blue, the clones are positively marked by expression of GFP (green). Scale bars: 50 *μ*m. Insets in top right corners show magnified images from the area indicated by dashed boxes. **B)** Violin plot shows averages of clone sizes for each wing disc (log2 scale), lines indicate average of each wing disc, and the violin shows kernel density estimate for the distribution of the average values. Asterisks indicate significant differences in the distributions (one-sided Mann-Whitney U test).

We and others have shown previously that loss of responsiveness of Myc to upstream oncogenic regulation can prevent tumorigenesis in animal models (*8*, *10*, *11*). The role of Myc target genes in tumorigenesis is less well understood. For example, decrease in expression of ribosomal proteins has been reported to have both positive (*48*) and negative (*49*) impact on tumorigenesis. Furthermore, loss of function of ribosomal proteins leads to decreased overall growth, and is linked to several inherited diseases (*50*). In our analyses, preventing the regulation of individual ribosomal proteins by Myc had very minor effects, suggesting that the effect of ribosomal protein haploinsufficiency on tumorigenesis is not necessarily specific to Myc, and could be instead caused by a more general limitation of growth. By contrast, we find here that blocking the ability of Myc to upregulate ribosome biogenesis abrogates its ability to drive growth. We show here that despite the complexity of the Myc regulated network, it is possible to specifically prevent regulation of a single Myc target gene in such a way that development proceeds largely normally, yet the ability of Myc to drive excessive growth is abrogated. This raises the possibility that Myc-driven growth could be specifically blocked by pharmacological agents, without severe side-effects. Furthermore, as ribosome biogenesis proceeds in multiple steps, it is possible that several enzymes in this pathway could be inhibited concomitantly to hinder the development of drug resistance.

## Materials and Methods

### Determining MYC/dMyc target genes conserved between human and *Drosophila melanogaster*

The list of MYC/dMyc targets conserved between human and Drosophila is based on fulfilling at least 3 out of 5 criteria in human (MYC binding (1), expression change upon CTNNB1 knockdown in microarray (2) or RNAseq (3) experiments, expression change upon MAX knockdown in microarray (4) or RNAseq (5) experiments) and 3 out of 6 of criteria in Drosophila melanogaster (dMyc (1) or Max (2) binding, expression change upon dMyc knockdown in microarray(3) or RNAseq (4) experiments, expression change upon Max knockdown in microarray (5) or RNAseq (6) experiments). The level of MYC knockdown in human was not sufficiently efficient to be included in the experiments within the criteria list, and with hence the values from CTNNB1 knockdown were used as the expression MYC dropped to below 5% of that in the control sample in RNAseq.

In human, binding within 1 kb from TSS was observed from ChIPexo experiments in LoVo human colon cancer cell line using an antibody against MYC (*51*) ChIP-exo data from (*51*) with ENA accession code: PRJEB9477) with peaks called by GPS (*52*). Microarray and RNAseq results in human where from GP5d colon cancer cell line, where genes with log2 fold changes > 1.5 upon CTNNB1 or MAX RNAi when compared to control samples were included. Affymetrix data was normalized with R package ‘affy’ using method ‘quantiles’ and annotated to Homo sapiens GRCh37.74 and only genes with normalized average expression among samples above 2 were considered. RNAseq results were analyzed by tophat2 and cuffdiff, mapped and annotated to iGenomes_hg19: UCSC, and only genes with expression above 5 FPKM in at least one sample out of 4 samples in the sequencing set were considered. Fastq and .CEL files will be made available in ArrayExpress.

Binding within 1 kb from TSS was observed from ChIPseq experiments in Drosophila S2 cells using antibodies against Myc or Max (Ref. (*16*); ArrayExpress accession code: E-MTAB-1648) and accepting peaks that have p values < 0.05 and fold changes > 1.5 to nonspecific antibody (IgG) when analyzed as described in (*53*).

Microarray and RNAseq results in Drosophila were based on data from (*53*)(*16*); ArrayExpress accession codes E-MTAB-453 and E-MTAB-1364 respectively(*53*), where genes with log2 fold changes > 1.5 upon dMyc or Max RNAi when compared to control samples (RNAi against GFP) in S2 cells were included. Additionally, in RNAseq, the number of reads for the gene had to be at least 1/100000 when sum of reads/sample equals 1, in at least one sample of GFP/dMyc/Max, and above 100 reads in at least one sample of all RNAseq samples (*16*). Orthologs were derived from Biomart using Ensembl genes 75 with Homo sapiens genes (GRCh37.p13) to Drosophila melanogaster genes (BDGP5).

### RNA-seq on wing imaginal discs

Conditional expression of UAS constructs in the wing pouch was achieved by using the nub-Gal4 and the TARGET system (*54*). Eggs were collected in bottles for 10h at 18°C, and then incubated for 4 days at 18°C (permissive temperature for Gal80TS), and finally transferred for 2 days to 29°C (restrictive temperature for Gal80TS) to induce expression of the UAS transgenes. For each replicate about 60 wing discs were dissected from wandering larvae and immediately transferred to the lysis buffer (Buffer RLT, Qiagen) supplemented with β-mercaptoethanol (1:100) and stored at −80°C until further processing.

Total RNA from each sample was isolated using the RNasey mini kit (Qiagen) according to manufacturer’s instruction and subsequently quantified with the Qubit 3.0 Fluorometer and the Qubit RNA BR Assay Kit (Thermo Fisher). RNA-seq libraries were generated with the Ovation Drosophila RNA-Seq System (NuGEN) which requires 10 and 100 ng of input RNA. Sequencing was performed using HiSeq4000 with 75 bp paired-end reads. STAR aligner mapped (BDGP5, Ensembl75 annotation) RNA-seq reads were analysed by DESeq2 (with default parameters and including all genes).

### Fly stocks

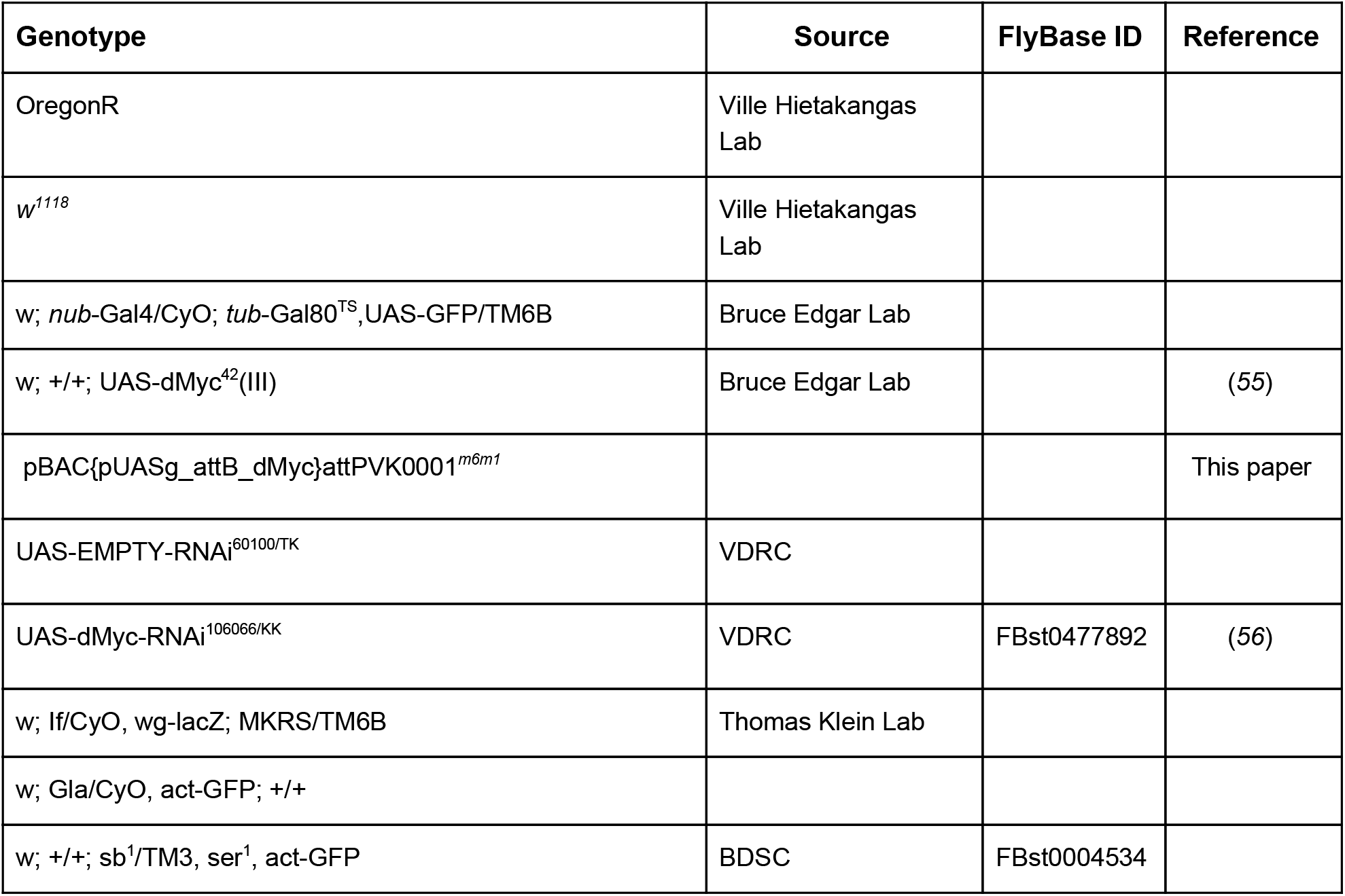
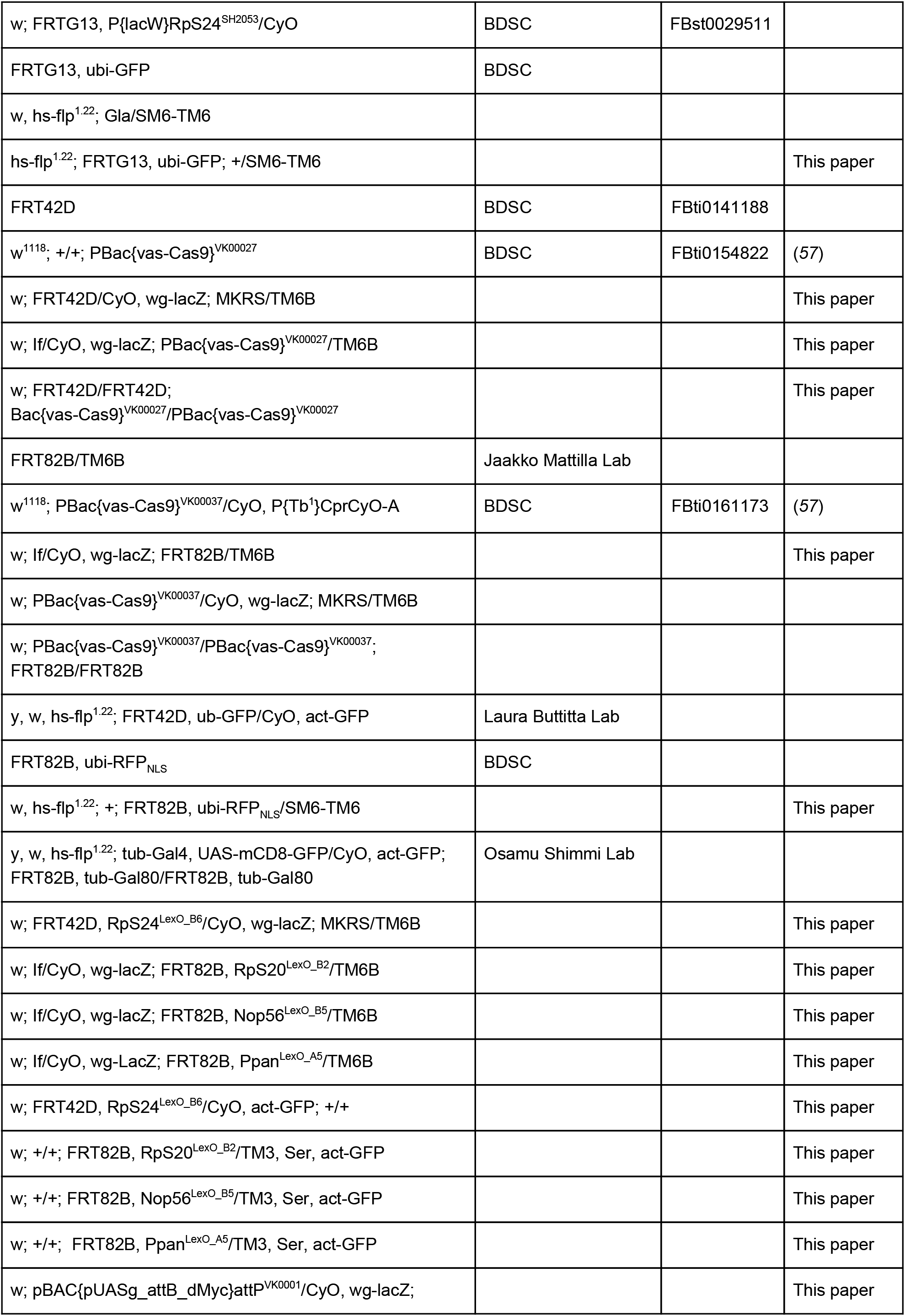
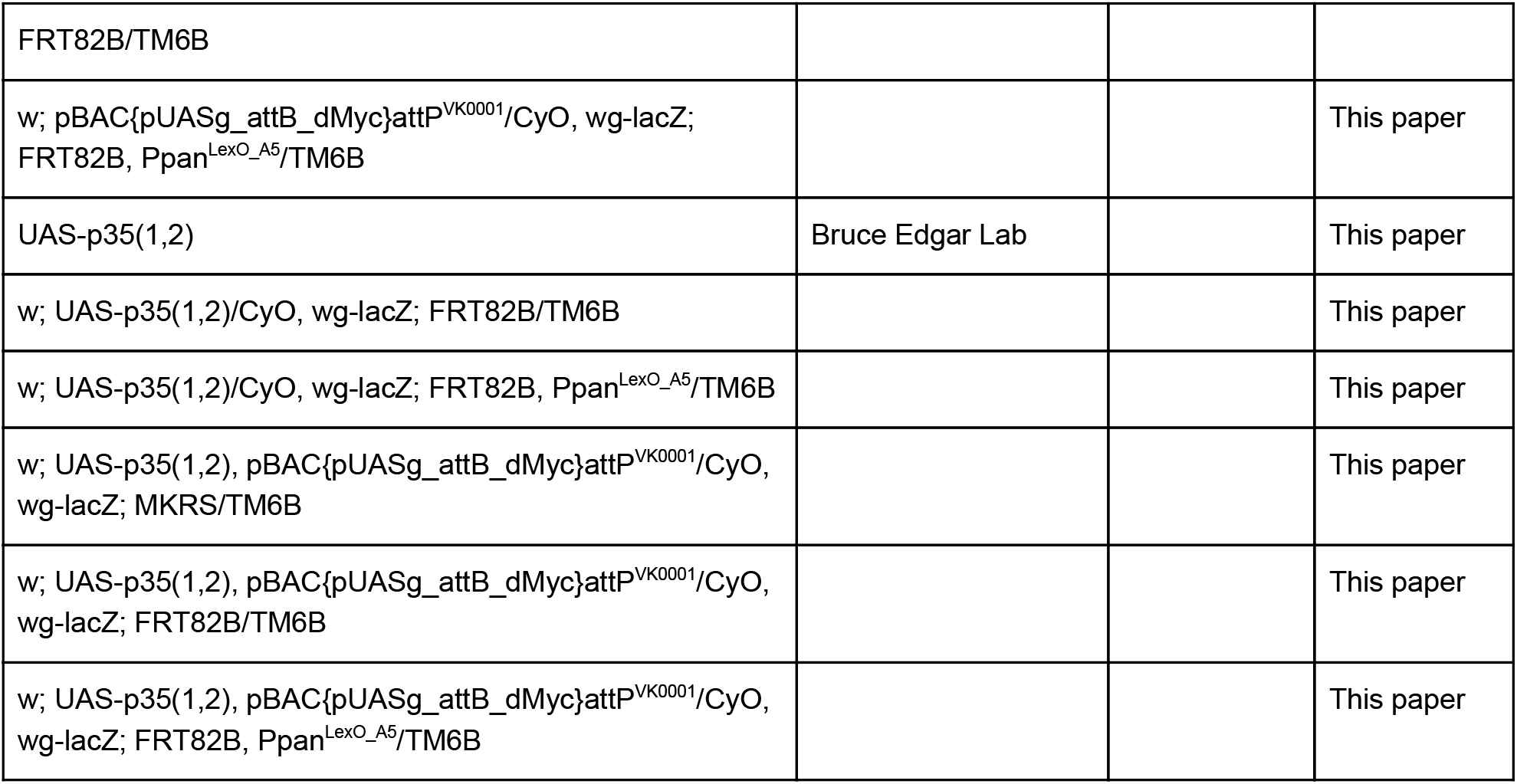

### CRISPR/Cas9-mediated gene editing in *Drosophila melanogaster*

Guide-RNAs were designed with the CRISPOR gRNA selection webtool (http://crispor.org) (*58*) and cloned into pCFD3: U6:3-gRNA as described by Port et al. (*59*).

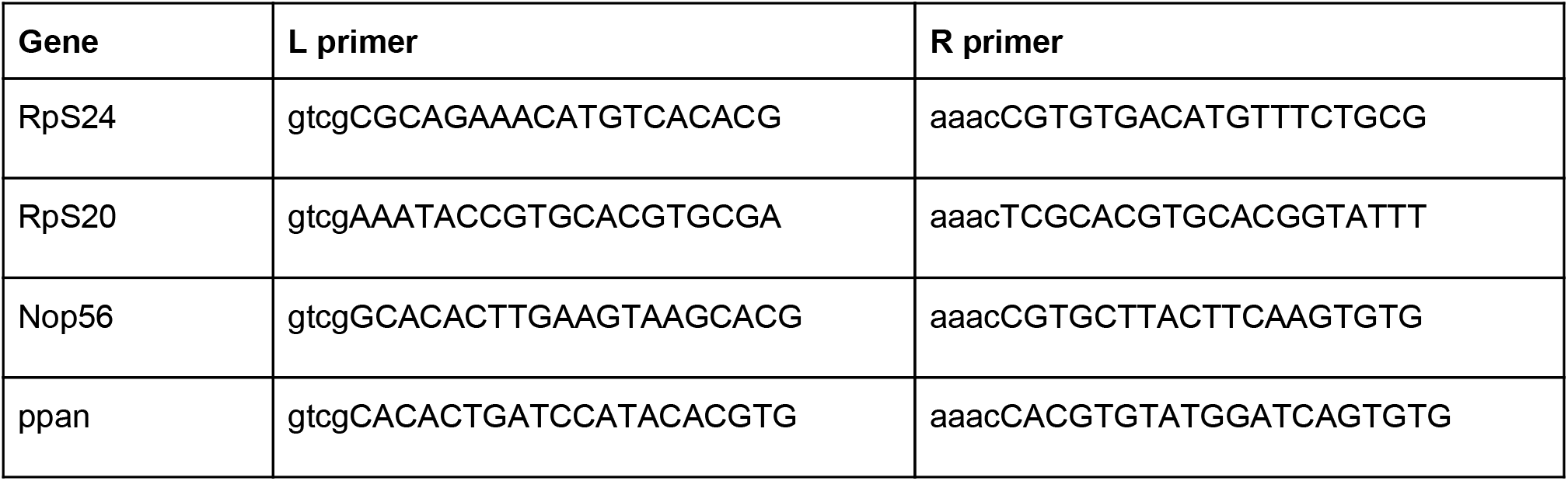

HR donor vectors containing the LexO sequence (CTGTATATATATACAG) in the position of the E-box as well as 1 kb homology arms (*60*) were created by gene synthesis (GenScript) and cloned into pUC57. Plasmids carrying gRNA and HR-donor were co-injected (GenetiVision) into *w; FRT42D/FRT42D; PBac{vas-Cas9}^VK00027^/PBac{vas-Cas9}^VK00027^* or *w; PBac{vas-Cas9}^VK00037^/PBac{vas-Cas9}^VK00037^; FRT82B/FRT82B*.

For genotyping (also see http://www.CRISPRflydesign.org/t7-endo-i-assay/) the resulting G0 flies were crossed twice with *w^1118^, If/CyO, wg-lacZ; MKRS/TM6B.* The founder males of the second cross were transferred to 96-well plates and lysed with *microLYSIS-PLUS* (Cambio) according to the manufacturer’s protocol. 1μl of each lysate was used as a PCR template to amplify 600-800 bp fragments around the edited E-box using the *Q5 High-Fidelity 2X Master Mix* (NEB). The resulting PCR products were purified in 96-well plates using the *AmpliClean DNA Cleanup Kit* (Nimagen) according to the manufacturer’s instructions. 17μl of the purified PCR products were then supplemented with 2μl NEBuffer2 and hybridized in a thermocycler using the following program : 5min, 95°C; ramp down to 85°C at −2°C/s; ramp down to 25°C at −0.1°C/s; hold at 4°C. Afterwards, 1 μl (10U) T7 endonuclease (NEB) were added to the hybridized PCR products and incubated for 15min at 37°C. The reaction was then stopped by adding 2μl of 0.25M EDTA and analyzed with the *Fragment Analyzer 5200* (Agilent) capillary electrophoresis instrument and the *CRISPR Discovery Gel Kit* (Agilent). To verify that the E-box was successfully replaced by the LexO site, the uncut PCR products of animals that showed a clear band-shift a were subcloned into *pCR-Blunt II-TOPO* (Thermo Fisher) and sequenced with SP6 primers.

Males from positive candidates were crossed twice against either *w; Gla/CyO, act-GFP; +/+* or *w; +/+; sb^1^/TM3, ser^1^, act-GFP*. Approximately 60 homozygous males from each E-box deleted fly line were disrupted in lysis-buffer (Buffer AL, Qiagen) using a motorized microtube homogenizer. Genomic DNA was isolated with the *DNeasy Blood & Tissue kit* (Qiagen) according to manufacturer’s instructions, with the exception that the gDNA was eluted in Low TE buffer (10 mM Tris-Cl, pH 8.0, 0.1 mM EDTA, Invitrogen). The concentration was measured with the *Qubit 3.0 Fluorometer* and the *Qubit dsDNA HS Assay Kit* (Thermo Fisher), and adjusted to a final concentration of 10 ng/μl. The region around the E-Box was amplified with primers outside of the homology regions using *Kappa HiFi HotStart ReadyMix* (Roche) and cloned into *pCR-Blunt II-TOPO* (Thermo Fisher). Correct insertion of the homology arms was then verified by Sanger sequencing. To rule out complex genomic rearrangements, the lines were also subjected to whole-genome sequencing (see below).

### Whole genome sequencing of edited fly lines

To confirm that the mutations affected the correct locus in the intended way, and were not the result of a complex genomic rearrangement, a DNA library was generated from 300 ng of genomic DNA using Kapa HyperPlus library preparation kit (Roche) according to manufacturers’ instructions. The library was then subjected to whole-genome sequencing using Illumina HiSeq4000 with 150 bp paired-end reads. The reads were mapped to *Drosophila* genome (Dmel_Release_6) and the correct structure of the locus was then confirmed manually. Four out of five lines had the expected genomic structure, whereas one line that appeared to be correctly targeted by PCR and Sanger sequencing had a genomic duplication that was detected by whole-genome sequencing. This line, RpL24^wt^:RpL24^Ebox−^, containing a tandem duplication of wt and E-box null RpL24 was therefore excluded from our study.

### Growth assays

To determine the pupariation timing of the E-box-deleted flies, eggs were collected for 4h at 37°C on apple juice agar plates. After 24h incubation at 37°C, the hatched larvae were manually genotyped based on GFP expression and for each genotype 3-5 replicates (vials) were prepared that contained 8-25 larvae. The number of pupae was quantified every 8h, starting at 94-100h after egg deposition.

To determine the emergence timing of the E-box-deleted flies, eggs were collected in bottles containing standard fly food for 4h at 37°C. The number of hatched flies was quantified every 8h, starting at 164-168h after egg deposition. The hatched flies were categorized by sex and then genotyped with help of the morphological markers of the respective balancer chromosomes.

### Mosaic analysis in *Drosophila* wing imaginal discs

Clonal analysis in wing discs was performed as described in (*61*). Briefly, eggs were collected for 1-4h at 25°C and incubated for another 48 h at 25°C when the flp recombinase was induced by a 1h heat-shock at 37°C. Imaginal discs were analysed when larvae had reached the wandering stage (about 3 days after clone induction).

### Immunohistochemistry

Imaginal discs were dissected in PBS and fixed for 30 min at 25°C in 4% paraformaldehyde/PBS. Primary antibodies: mouse anti-dMyc P4C4-B10 (1:10, The Developmental Studies Hybridoma Bank at the University of Iowa) and Rabbit anti-cleaved *Drosophila* Dcp-1 (Asp216) (1:100; Cell Signaling Technologies). Secondary antibodies were used at a dilution of 1:500. DNA was visualized with Hoechst 33342 (1:1000 of 10mg/ml stock solution, Sigma Aldrich).

### Confocal Microscopy and image analysis

Z-stacks (0,68 μm) of imaginal discs were taken using a Leica SP8 confocal microscope using 20X lens at 0.9x magnification. The images were processed with Fiji using the rolling ball algorithm for background subtraction. Each image represents a maximum z-projection of 5 slices. The imaginal discs and clone boundaries were then manually identified, and their areas quantified using. For each disc, the fraction of GFP positive clones were then measured, and the numbers plotted to **Figure 3**. The violin plot was generated using Python packages Matplotlib and Seaborn, function “catplot”, using options kind=“violin”, inner=“stick”, and palette=“pastel”. Statistical significance was assessed using Mann-Whitney test from the package Scipy, function “mannwhitneyu” using default settings (with continuity correction, one-sided test). Brackets and asterisks were added manually, by using package Matplotlib pyplot, functions “plot” and “text”.

### Scanning electron microscopy and image analysis

Females of the indicated genotypes were fixed overnight at 4°C with rocking in 2% glutaraldehyde in PBS. After dehydration in a series of ethanol dilutions, the dried flies were coated with platinum using a Q150T turbo-pumped sputter coater (Quorum) under 0.3 bar argon atmosphere for 25 seconds (approx. 5 nm coat). Scanning electron microscopy analysis was performed using a FEI Quanta 250 FEG instrument with the following settings (HV: 5.00 kV, pressure 4.07 × 10^−3^ Pa, dwell: 3 μs, spot: 3.5, WD: 11.2 mm, magnification: 250x). Scutellar bristle lengths (**Figure 2**) were quantified using the Microscopy Image Browser developed by the Electron Microscopy Unit at the Institute of Biotechnology, University of Helsinki. Scutellar bristles were colorized using the layer mask tool in Photoshop CC 2014 (Adobe). P-values were calculated with Prism 8.0 (Graphpad Software) using Welch’s T-test.

## Acknowledgments

We thank Drs. Laura Buttitta, Bruce Edgar, Ville Hietakangas, Jaakko Mattila, and Osamu Shimmi, as well as the Vienna and Bloomington Stock Centers for *Drosophila* stocks, and Drs. Ville Hietakangas and Minna Taipale for critical review of the manuscript. Anu Luoto, Kaisu Jussila and Katariina Sarin for technical assistance. We also thank the *Drosophila* (Hi-Fly), Light (BIU) and Electron Microscopy (EMBI) facilities at the University of Helsinki and the Next Generation Sequencing Services Institute for Molecular Medicine Finland (FIMM).

## Notes

### Competing Interest Statement

The authors have declared no competing interest.

